# Adolescent Binge Drinking is Associated with Accelerated Decline of Gray Matter Volume

**DOI:** 10.1101/2021.04.13.439749

**Authors:** M.A. Infante, Y. Zhang, T. Brumback, S.A. Brown, I.M. Colrain, F.C. Baker, D.B. Clark, D. Goldston, B.J. Nagel, K.B. Nooner, Q. Zhao, K.M. Pohl, E.V. Sullivan, A. Pfefferbaum, S.F. Tapert, W.K. Thompson

## Abstract

The age- and time-dependent effects of binge-drinking on adolescent brain development have not been well characterized even though binge drinking is a health crisis among adolescents. The impact of binge drinking on gray matter volume development was examined using longitudinal data from the National Consortium on Alcohol and NeuroDevelopment in Adolescence (NCANDA). Non-binge drinkers (n=177) were matched to binge drinkers (n=164) on potential confounders. Number of binge drinking episodes in the past year was linked to decreased volumes for total gray matter, frontal, parietal, temporal, and occipital lobes (*p*s<.001). Interactions of binge drinking episodes and age demonstrated stronger effects in younger subjects for total gray matter, frontal, temporal, and occipital lobes (*p*s<.001). Subsequent models included binge drinking coded in multiple ways. Models sensitive to number of episodes and temporal proximity to outcomes provided the best fits. Declines in gray matter volume association with binge drinking are potentially related to changes in cognition frequently reported among binge drinking adolescents. Results underscore the potential importance of delaying initiation of binge drinking and provide evidence for a dose-response relationship of binge drinking to gray matter decline. Temporally proximal binge drinking was associated more strongly with gray matter decline, suggesting the potential for recovery.

---

Adolescence is a critical developmental stage when experimentation with alcohol and other substances is often initiated. Alcohol remains the most widely used substance during adolescence with 19%, 38%, and 52% reporting past-year alcohol use by 8th, 10^th^, and 12th grade respectively (Johnston et al. 2020). Although adolescents generally consume alcohol with less frequency than adults, among those who report drinking the majority engages in binge drinking (Substance Abuse and Mental Health Services Administration 2020), which is defined by the National Institute on Alcohol Abuse and Alcoholism (NIAAA) as a pattern of drinking alcohol that brings the blood alcohol concentration (BAC) to 0.08g/dl or higher (2004). Findings from the most recent Monitoring the Future survey show 14.4% of adolescents in the 12^th^ grade had engaged in binge drinking in the past two weeks and data from the 2019 National Survey on Drug Use and Health shows 4.9% of adolescents aged 12-17 had engaged in past month binge drinking. Binge drinking has been identified as particularly risky for adolescents, with far more severe consequences compared to adults, including deviation from normal brain development, (Zhao et al. 2020) structural and functional brain alterations, (Jones et al. 2018) neuropsychological deficits, (Carbia et al. 2018) and increased risk for developing alcohol as well as other substance use disorders (Chassin et al. 2002). The burden of excessive alcohol use alone on US society is estimated to be over 223 Billion a year (Bouchery et al. 2011).

During adolescence the brain goes through significant structural and organizational changes (Gogtay et al. 2004; Shaw et al. 2008). Longitudinal structural magnetic resonance imaging (MRI) studies have shown region-specific variation of gray matter volume reduction from childhood through early adulthood, with sensorimotor and occipital brain regions maturing first, followed by limbic regions important for rewards and emotion, and higher order cortical association areas, including the prefrontal cortex maturing later in development (Gogtay et al. 2004; Sowell et al. 2004). The asynchronized development of brain regions, particularly the difference in timing of the development of the reward and control systems may explain why adolescents might be more prone to risk-taking behaviors (Crews et al. 2007; Casey et al. 2008). At the same time, given the extensive nature of neurodevelopment during this period, adolescents’ brains may be especially vulnerable to intoxicants such as alcohol.

Findings from cross-sectional studies have shown gray matter differences in binge drinking college-age adolescents compared to their non-drinking peers, including smaller volumes in the frontal lobe (Kvamme et al. 2016) and cerebellum (Lisdahl et al. 2013). In contrast, others have found larger cortical volumes in college-age binge drinkers compared to non-drinkers, including regions of the prefrontal cortex, anterior cingulate (Doallo et al. 2014) and ventral striatum (Howell et al. 2013). Greater gray matter density in the left middle frontal gyrus among college-age binge drinkers than non-drinkers was also reported in a more recent study (Sousa et al. 2017). Sex differences in the effect of binge drinking on brain structure have also been documented, perhaps related to sex-related differences in genetic vulnerabilities or neurotoxic sensitivities (Kvamme et al. 2016).

Longitudinal studies suggest an accelerated decline in cortical volumes may result from heavy alcohol use during adolescence (Squeglia et al. 2014; Squeglia et al. 2015; Meda et al. 2017; Pfefferbaum et al. 2018). A study by Squeglia et al. (2014) examined brain development before and after initiation of heavy drinking, showing alcohol-associated reduction in subcortical regions as well as inferior and middle temporal structures. In another study (Meda et al. 2017), subjects were categorized as heavy versus light drinkers based on number of binge drinking episodes in the previous 26 weeks, with subjects in the heavy binge drinking group showing an increased rate of gray matter decline, primarily in fronto-striatal regions. Using three waves of data from the National Consortium on Alcohol and NeuroDevelopment in Adolescence (NCANDA), Pfefferbaum et al. (2018) examined the impact of adolescent heavy drinking on brain trajectories of gray and white matter volumes. That study compared brain trajectories of those who at the two-year follow-up remained no/low drinkers versus those who had initiated moderate or heavy drinking by this time point. Results indicated smaller volumes of frontal, cingulate, and total gray matter in heavy drinkers compared with those in the no/low drinking group. As in Squeglia et al. (2014), subjects were classified based moderate or heavy drinking pattern as opposed to binge drinking status. Understanding the impact of binge drinking on adolescent brain development of specific brain regions has important implications given the association between neurobiological changes at the structural and functional level to cognitive and behavioral changes (Jones et al. 2018).

To date, the age- and time-dependent effects of binge-drinking on adolescent brain development have not been well characterized. Large-scale longitudinal studies with several years of data collection are needed to further understand the effects of binge drinking, including its proximal and longer-term effects on the brain. To begin to address these issues, the present study expands on the Pfefferbaum et al.(2018) analysis in three ways. <u>First</u>, we assessed the relationship of adolescent binge drinking with gray matter volume trajectories across all five extant waves of longitudinal data from the NCANDA study, encompassing a longer time period, broader age ranges, and a larger variation in drinking behaviors than in prior reports using NCANDA data. <u>Second</u>, we leveraged the NCANDA cohort sequential design to assess whether strength of any associations of binge drinking with gray matter volume trajectories varied by baseline age. <u>Third</u>, we examined whether models that are sensitive to number of binge drinking episodes (“dose”) and temporal proximity of binge drinking to outcomes would explain more variation than models that incorporate binge drinking in other ways (as discrete indicators and/or as cumulative measures). The NCANDA study’s large sample size, number of longitudinal assessments and cohort sequential design thus allow for an unprecedented investigation into the nature of the association of adolescent alcohol use and subsequent trajectories of cortical gray matter volume development across adolescence and into young adulthood.

## Materials and Methods

### Subjects

Data came from the National Consortium on Alcohol and Neurodevelopment in Adolescence (NCANDA). The NCANDA cohort consists of 831 12 to 21 year-olds enrolled across five sites (Duke University, University of Pittsburgh Medical Center, Oregon Health & Science University, University of California San Diego, and SRI International) and followed for 5 years (Brown et al. 2015). Subjects were recruited through local schools and targeted catchment-area calling. The study follows a cohort sequential design, which recruited youth in three age bands (12-14, 15-17, and 18-21 years), enabling examination of a broad developmental window due to between-subject variation in baseline age (“age cohort”). Prior to study entry, most of the sample had not engaged in binge drinking and ~15% (n=121) had at least one prior binge drinking episode. Youth at risk for heavy drinking (i.e., early experimentation with alcohol, family history of substance use disorder, externalizing or internalizing symptoms) were overrecruited to comprise 50% of the sample.

Exclusion criteria included lack of English fluency, MRI contraindications, serious medical conditions (e.g., traumatic brain injury with loss of consciousness >30 minutes), non-correctable sensory impairments, current serious Axis I psychiatric disorder that may influence study completion (e.g., psychosis), and early developmental problems (e.g., known exposure to prenatal alcohol or other drugs) (Brown et al. 2015). The research protocol was approved by the Institutional Review Board at each site. At each time point, subjects (or the parent or legal guardian for minors) provided written informed consent and written informed assent was obtained from minors. Subjects and their parents were compensated for participation.

After informed consent, at baseline and each follow-up year subjects completed a comprehensive assessment of substance use, psychiatric symptoms, functioning in major life domains, neuropsychological testing, and neuroimaging (Brown et al. 2015). This included the Customary Drinking and Drug use Record (CDDR) (Brown et al. 1998) to characterize past and current alcohol and other substance use, reported on past-year use frequency, maximum number of drinks in a drinking episode, and number of binge drinking episodes (i.e., 5 or more drinks for males or 4 or more drinks for females on an occasion). The number of binge drinking episodes in the prior year at each annual follow-up assessment was the primary (longitudinal) independent variable of interest.

Subjects who reported having had at least one lifetime binge episode at baseline (n=121) were excluded from analysis. Because the focus of this paper is longer-term temporal variation in binge drinking, we excluded subjects who had not completed all five (baseline and four annual follow-up) visits to allow sufficient within-subject assessments of binge drinking to characterize its association with concurrent and subsequent gray matter volumes. This resulted in a final pool of n=341 subjects for analyses. Of these, n=177 (52%) reported never having had a binge drink episode over the five visits and n=164 (48%) reported binge drinking at least once during the study period subsequent to baseline. Subject characteristics are reported in Table 1.

**Table 1:** Subject characteristics (% or mean) for subjects with at least one binge-drinking episode (Binge Drinkers) vs. those with no binge-drinking episodes (Controls) before and after matching on baseline covariates.

|  | **Binge Drinkers**<br>**(n=164)** | **Before matching**<br>**Controls (n=177)** | **Post matching**<br>**Controls (n=78)** |
| --- | --- | --- | --- |
| Sex (% male) | 52.44 | 49.72 | 52.44 |
| Baseline Visit Age in years (sd) | 16.53 (1.9) | 15.05 (2.4) | 16.23 (2.2) |
| Site |  |  |  |
| Site A (%) | 7.93 | 13.00 | 20.72 |
| Site B (%) | 10.37 | 7.91 | 7.93 |
| Site C (%) | 14.63 | 22.03 | 12.20 |
| Site D (%) | 25.00 | 22.03 | 22.56 |
| Site E (%) | 42.07 | 35.03 | 36.59 |
| Race |  |  |  |
| Asian (%) | 7.93 | 5.65 | 10.37 |
| Caucasian (%) | 80.49 | 70.06 | 80.49 |
| Other (%) | 4.89 | 6.78 | 4.29 |
| Socioeconomic status<sup>+</sup>(sd) | 16.95 (2.3) | 16.17 (2.7) | 16.64 (2.3) |
| Baseline ICV<sup>++</sup> (sd) | 0.13 (1.0) | -0.12 (0.99) | 0.10 (0.92) |

<sup>+</sup> Highest level of education achieved by either parent at baseline
<sup>++</sup> Total intracranial volume standardized to have zero mean and unit variance

Binge drinkers (those with at least one binge drinking episode across all five visits) were matched with non-binge drinkers (subjects with no binge drinking episodes at any visit) on potential confounders, including sex, age, race/ethnicity, highest level of education completed by either parent, site and baseline (pre-binge drinking) intracranial volume (ICV). Matching was performed by first computing a propensity score with binge-drinker (yes/no) as the dependent variable and each of these matching variables as covariates in a multiple logistic regression. The non-binge drinker with the closest propensity score to a given binge drinker was selected as the matched control. Because the pool of non-binge drinkers to select from was only slightly larger than the number of binge drinkers, we performed matching *with replacement* to improve the post-match balance on these potential confounders. Table 1 displays the pre- and post-match summaries of potential confounders of binge and non-binge drinkers, generally demonstrating improved balance for potential confounders, with the sole exception of site. (All matching variables were also included in subsequent analyses to control for any residual imbalances.)

### MRI Data Acquisition and Analysis

A high-resolution structural MRI protocol and the use of 3T scanners was consistent across sites. Three sites (UCSD, SRI, and Duke University) used GE MR750 and two sites (UPMC and OHSU) used Siemens TIM-Trio scanners. T1-weighted and T2-weighted images were acquired in the sagittal plane. Descriptions of the structural MRIs can be found elsewhere. (Pfefferbaum et al. 2016; Pfefferbaum et al. 2018). The imaging data processing and selection of regions for this study is based (Pfefferbaum et al. 2018) and details can be found there. Briefly, following skull-stripping, FreeSurfer software (version 5.3, http://surfer.nmr.mgh.harvard.edu) was then used for the volumetric parcellation and segmentation to estimate total gray matter volume as well as volumes of frontal, temporal, parietal, occipital, cingulate and insular cortices from the Desikan-Killiany cortical atlas (Desikan et al. 2006). The SRI24 atlas (Rohlfing et al. 2010; Rohlfing et al. 2014) was used to measure ICV volume. Volumes (expressed in mm^3^) were combined across hemispheres to produce one value for each ROI.

### Statistical Analyses

Primary analyses consisted of seven separate linear mixed-effects models (LMEs), one for each gray matter volume (GMV) of interest: total GMV, frontal lobe, parietal lobe, temporal lobe, occipital lobe, insula, and cingulate. An LME was fit with each of the seven GMVs (standardized to have zero mean and unit standard deviation) as the dependent variable in turn. The primary independent variable of interest was number of binge drinking episodes in the prior year (*binge_ij_*) for the *i*th subject at the *j*th visit (*j* = 1,…,5). Because of the long tail in number of episodes (see Figure 1), *binge_ij_* was log transformed (after adding one to avoid taking the log of zero) and then mean centered at zero. Age was entered into the LME as two terms to reflect the cohort sequential aspect of the NCANDA study design. <u>Between-subject</u> variation in ages was captured by the mean age of the subject across the five visits (*age_m,i_*). <u>Within-subject</u> change in age (*age_d,ij_*) was captured by subtracting *age_m,i_* from the *i*th subject’s age at visit *j*. Each of these terms (*binge_ij_*, *age_m,i_*, and *age_d,ij_*) was entered into LMEs along with their two-way interactions. Random intercepts were included for subject and family. Variables used in matching were also included as fixed effects to control for any residual imbalances after matching. We assessed the significance of binge drinking on gray matter volumes by fitting the same model without *binge_ij_* and using a Likelihood Ratio Test (LRT). All reported p-values were two-sided and Bonferroni-corrected for seven comparisons (*p*=.007).

**Figure 1.**
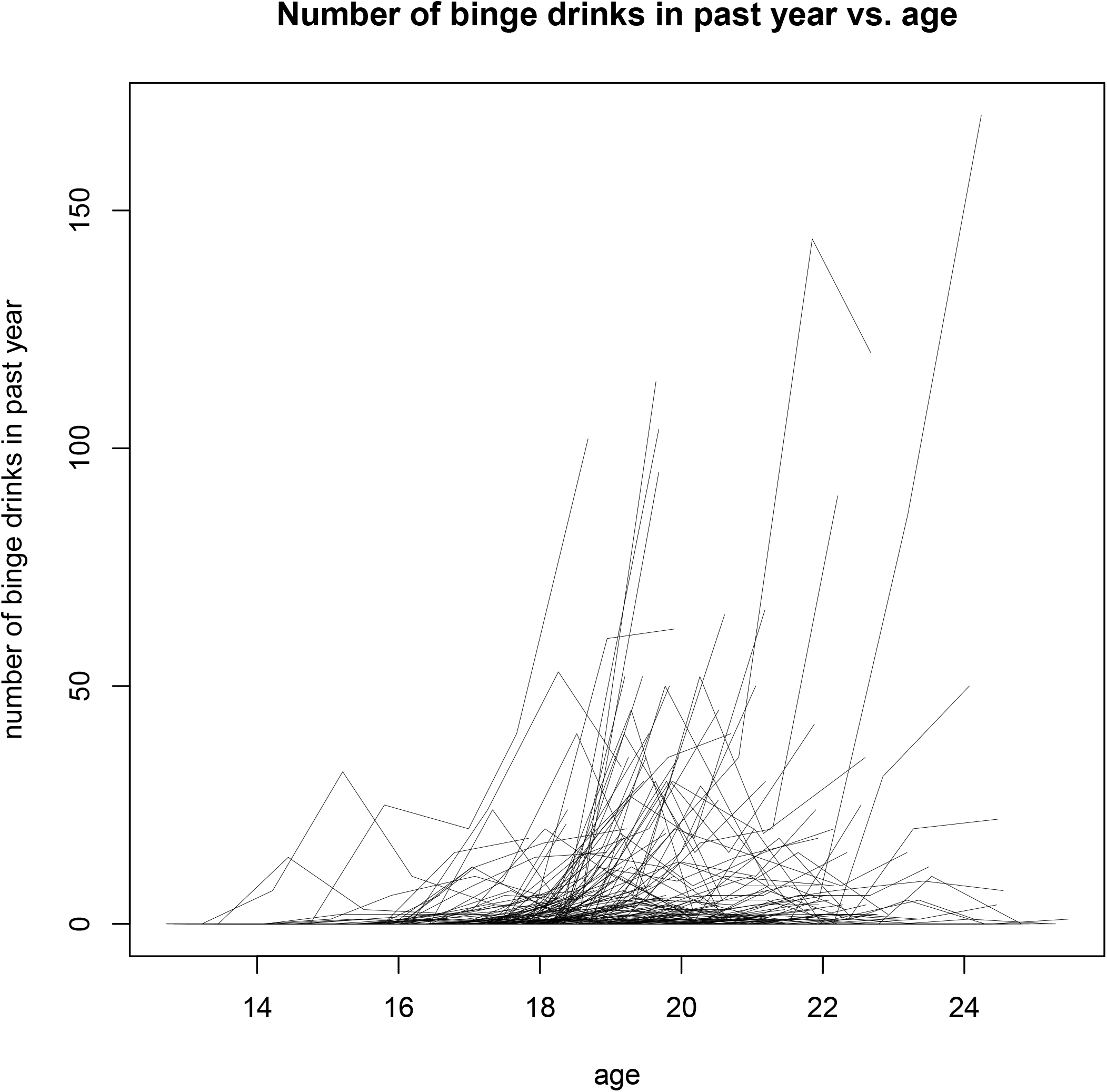
Number of binge drinking episodes in the prior year by age (N=242).

We then performed a second set of analyses by comparing model fits of coding binge drinking in several ways: M0) a base model without any binge drinking variable; M1) number of binge drinking episodes in the prior year (*binge_ij_*,), identical to the primary analyses; M2) cumulative number of binge drinking episodes (*binge_cum_ij_*), equal to the log-transformed sum of the number of current and past binge drinking episodes; M3) *binge_cat_ij_*, equal to one if the subject had one or more binge-drinking episodes in the last year and zero otherwise; M4) current or past binge drinking (*binge_past_ij_*), equal to one if the subject had a binge drinking episodes at the current visit or at any past visit; M5) ever binge drank (*binge_ever_i_*) equal to one if the subject had at least one binge drink in any visit and zero otherwise; and M6) log-transformed total number of binge drinking episodes across all five visits (*binge_tot_i_*). These different ways of coding binge drinking were entered into separate LMEs (6 coding x 7 volumes = 42 total models) for each of the GMV’s as described above for *binge_ij_* and the fits were compared using the Akaike Information Criterion (AIC) (Akaike 1974). All statistical analyses were conducted using R Version 3.6.3. (R Core Team 2020).

## Results

Demographic characteristics of the sample at baseline (N=341) by binge drinking status before, 177 (51.9%) and after, 78 (32.2%) matching controls are presented in Table 1. After matching the final sample consisted of 242 subjects (164 binge drinkers [68.8%] with mean [SD] age 16.5 [1.89] years, 86 male [52.4%]; and 78 (32.2%) non-binge drinkers with mean [SD] age 16.23 [2.24] years, 41 male [52.5%]). Table 2 gives the distribution of binge drinking by age cohort for each yearly follow-up assessment. Figure 1 displays a spaghetti plot of binge-drinking trajectories across all five yearly assessments. Information on subjects’ cannabis and tobacco use can be found in Supplementary Materials eTable1.

**Table 2:** Mean number (range, sd) of binge drinking episodes in past year by follow-up year by age cohort for N = 164 subjects with at least one binge drinking episode over the NCANDA study period to date.

| Baseline Age Cohort<br>(Years) | Mean number of binge drinking episode in the past year (range, sd) |  |  |  |
| --- | --- | --- | --- | --- |
|  | Y1 | Y2 | Y3 | Y4 |
| Cohort 1 (12-14) | 1.5 ([0,25], 4.9) | 2.4 ([0,32], 6.5) | 3.1 ([0,40], 7.3) | 9.0 ([0,102], 17.3) |
| Cohort 2 (15-17) | 0.7 ([0,12], 1.7) | 4.4 ([0,45], 8.6) | 7.5 ([0,60], 12.2) | 15.5 ([0,114], 23.7) |
| Cohort 3 (18-21) | 3.7 ([0,20], 6.2) | 5.9 ([0,52], 10.6) | 12.7 ([0,144], 27.2) | 18.1 ([0,170], 36.4) |

In the first set of seven LMEs, the LRTs of models with *binge_ij_* vs. models without were nominally significant at the alpha = 0.05 level for all seven GMV dependent variables. After a Bonferroni correction, all were still significant except for cingulate and insula. LME fixed-effect regression coefficient estimates (along with their Wald test p-values and 95% confidence intervals) as well as the overall model LRT statistics and p-values are presented in Table 3. Spaghetti plots of trajectories and model fits are displayed in Figure 2. All GMVs were smaller as a function of both, *age_m,i_* and *age_d,ij_*. In all seven LMEs, the main effects of *binge_ij_* were negative, the two-way interactions of *binge_ij_* × *age_m,i_* were significant for all regions except the insula after Bonferroni correction and were consistently positive for all models, indicating attenuated association of volumes with number of binge drinking episodes in older subjects.

**Table 3:**
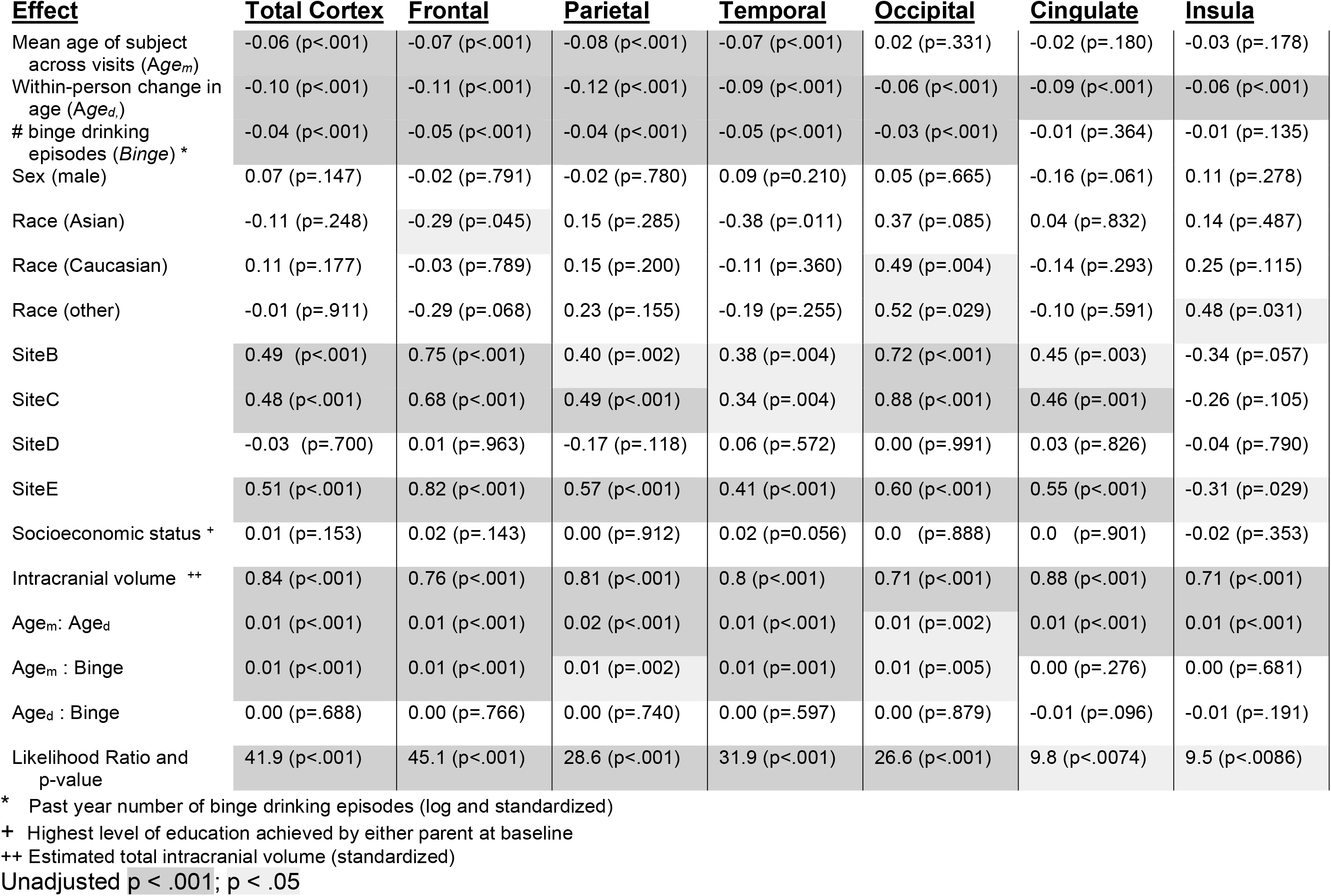
Linear mixed effects (LME) coefficients and p-values for each gray matter region of interest.

| **Effect** | **Total Cortex** | **Frontal** | **Parietal** | **Temporal** | **Occipital** | **Cingulate** | **Insula** |
| --- | --- | --- | --- | --- | --- | --- | --- |
| Mean age of subject across visits ($Age_m$) | -0.06 (p<.001) | -0.07 (p<.001) | -0.08 (p<.001) | -0.07 (p<.001) | 0.02 (p=.331) | -0.02 (p=.180) | -0.03 (p=.178) |
| Within-person change in age ($Age_d$) | -0.10 (p<.001) | -0.11 (p<.001) | -0.12 (p<.001) | -0.09 (p<.001) | -0.06 (p<.001) | -0.09 (p<.001) | -0.06 (p<.001) |
| # binge drinking episodes ($Binge$) * | -0.04 (p<.001) | -0.05 (p<.001) | -0.04 (p<.001) | -0.05 (p<.001) | -0.03 (p<.001) | -0.01 (p=.364) | -0.01 (p=.135) |
| Sex (male) | 0.07 (p=.147) | -0.02 (p=.791) | -0.02 (p=.780) | 0.09 (p=0.210) | 0.05 (p=.665) | -0.16 (p=.061) | 0.11 (p=.278) |
| Race (Asian) | -0.11 (p=.248) | -0.29 (p=.045) | 0.15 (p=.285) | -0.38 (p=.011) | 0.37 (p=.085) | 0.04 (p=.832) | 0.14 (p=.487) |
| Race (Caucasian) | 0.11 (p=.177) | -0.03 (p=.789) | 0.15 (p=.200) | -0.11 (p=.360) | 0.49 (p=.004) | -0.14 (p=.293) | 0.25 (p=.115) |
| Race (other) | -0.01 (p=.911) | -0.29 (p=.068) | 0.23 (p=.155) | -0.19 (p=.255) | 0.52 (p=.029) | -0.10 (p=.591) | 0.48 (p=.031) |
| SiteB | 0.49 (p<.001) | 0.75 (p<.001) | 0.40 (p=.002) | 0.38 (p=.004) | 0.72 (p<.001) | 0.45 (p=.003) | -0.34 (p=.057) |
| SiteC | 0.48 (p<.001) | 0.68 (p<.001) | 0.49 (p<.001) | 0.34 (p=.004) | 0.88 (p<.001) | 0.46 (p=.001) | -0.26 (p=.105) |
| SiteD | -0.03 (p=.700) | 0.01 (p=.963) | -0.17 (p=.118) | 0.06 (p=.572) | 0.00 (p=.991) | 0.03 (p=.826) | -0.04 (p=.790) |
| SiteE | 0.51 (p<.001) | 0.82 (p<.001) | 0.57 (p<.001) | 0.41 (p<.001) | 0.60 (p<.001) | 0.55 (p<.001) | -0.31 (p=.029) |
| Socioeconomic status + | 0.01 (p=.153) | 0.02 (p=.143) | 0.00 (p=.912) | 0.02 (p=0.056) | 0.0 (p=.888) | 0.0 (p=.901) | -0.02 (p=.353) |
| Intracranial volume ++ | 0.84 (p<.001) | 0.76 (p<.001) | 0.81 (p<.001) | 0.8 (p<.001) | 0.71 (p<.001) | 0.88 (p<.001) | 0.71 (p<.001) |
| $Age_m$: $Age_d$ | 0.01 (p<.001) | 0.01 (p<.001) | 0.02 (p<.001) | 0.01 (p<.001) | 0.01 (p=.002) | 0.01 (p<.001) | 0.01 (p<.001) |
| $Age_m$ : Binge | 0.01 (p<.001) | 0.01 (p<.001) | 0.01 (p=.002) | 0.01 (p=.001) | 0.01 (p=.005) | 0.00 (p=.276) | 0.00 (p=.681) |
| $Age_d$ : Binge | 0.00 (p=.688) | 0.00 (p=.766) | 0.00 (p=.740) | 0.00 (p=.597) | 0.00 (p=.879) | -0.01 (p=.096) | -0.01 (p=.191) |
| Likelihood Ratio and p-value | 41.9 (p<.001) | 45.1 (p<.001) | 28.6 (p<.001) | 31.9 (p<.001) | 26.6 (p<.001) | 9.8 (p<.0074) | 9.5 (p<.0086) |

\* Past year number of binge drinking episodes (log and standardized)
+ Highest level of education achieved by either parent at baseline
++ Estimated total intracranial volume (standardized)
Unadjusted p < .001; p < .05

**Figure 2.**
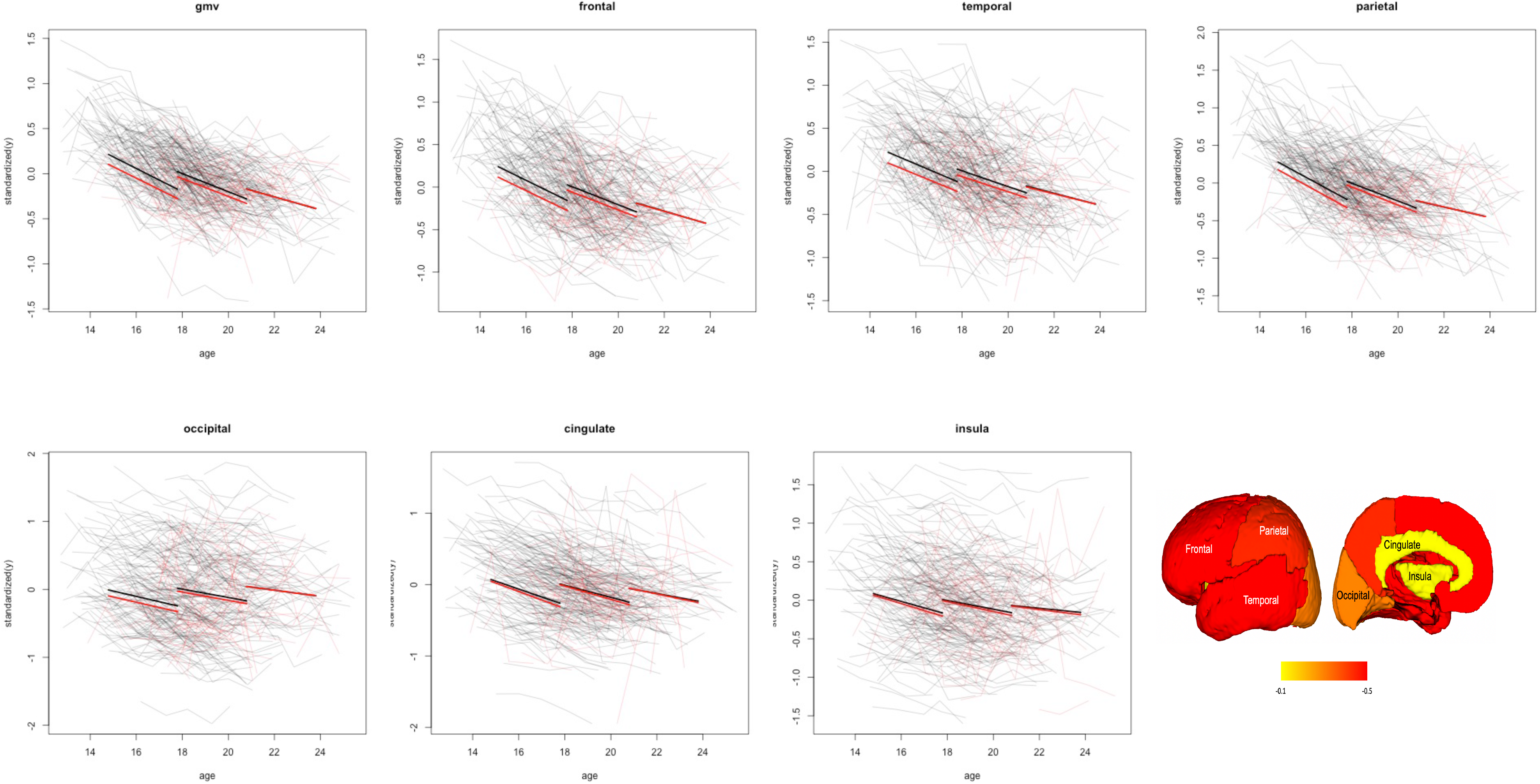
Spaghetti plots showing group differences between binge (n=164; shown in green) and matched control (n=78; shown in gray) subjects on brain developmental trajectories and model fits by age group categories.

We also examined the potential differential effects of binge drinking for males and females by including an interaction of sex with *binge_ij_* and testing these vs. models with no interaction using LRTs. None of the interactions were significant (unadjusted *ps* = 0.228 to 0.751).

The AIC results of the second set of LMEs are presented in Table 4. For each set of six models, the AIC for the base model M0 was subtracted to highlight the improvement of model fit of including binge drinking from the model with no binge drinking term: lower (here, more negative) AIC indicates better model fit than that for M0. For all seven gray matter volumes, all models including a binge drinking term (M1-M6) were better fitting than M0. Moreover, the temporally proximal dimensional model (M1) was the best fitting for all GMVs except for insula (which in any case had the weakest association with binge drinking which was not statistically significant after Bonferroni correction). The second-best fitting model was M2 with the exception of insula and cingulate. The worst fitting models were generally M4-M6. In sum, including both “dose” and temporal proximity were conjointly informative and strongly improved upon fits of models ignoring this information.

**Table 4:**
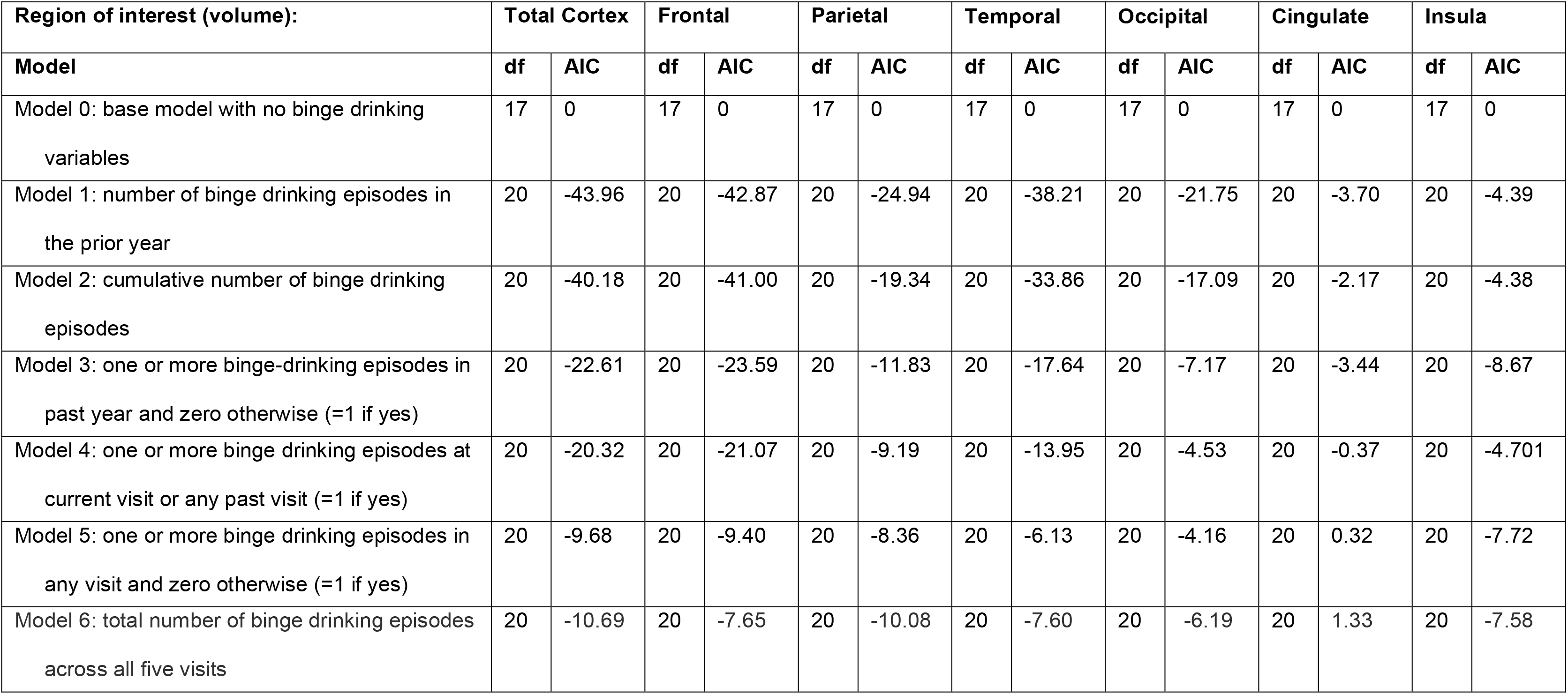
Akaike Information Criterion (AIC) of 6 model comparisons coding binge drinking in different ways.

## Discussion

The goal of this study was to build on existing research to advance our understanding of the age- and time-dependent effects of binge drinking on adolescent brain development. Using five-waves of longitudinal data from the cohort sequential NCANDA study, we examined the extent to which binge drinking disturbs the development of gray matter volumes. We specifically focused on binge drinking given the high rates of binge drinking among adolescents (Substance Abuse and Mental Health Services Administration 2020), preliminary evidence for the negative effects of binge drinking on the adolescent brain (Jones et al. 2018), and the limited longitudinal neuroimaging studies in this area. We found that binge drinking was associated with lower gray matter volumes. Specifically, our results revealed greater number of binge drinking episodes in the prior year was associated with decline in frontal, parietal, temporal, occipital and total gray matter volumes. Further, we found the association between gray matter volumes and binge drinking was attenuated among older subjects. Finally, findings from our comparative analyses revealed that models coding binge drinking as a dimensional, temporally proximal variable better fit the data versus models ignoring this information. Taken together, these results provide evidence for a dose-response relationship of binge drinking with reduced gray matter volumes in younger as opposed to older youths, as well as and evidence for time-dependent associations, with temporally proximal number of binge drinks more predictive of reduced gray matter than cumulative or total lifetime binge drinks.

Our results showing decreased gray matter volumes concurrent with binge drinking episodes are consistent with prior longitudinal studies on moderate to heavy drinking youth (Squeglia et al. 2014; Squeglia et al. 2015; Meda et al. 2017; Pfefferbaum et al. 2018). The association of binge drinking with lower gray matter volumes may be mediated by neurotoxic effects related to accelerated maturation in gray matter pruning, as suggested by Pfefferbaum et al. (2018). Alternatively, pre-existing differences may account for the association of brain developmental trajectories and binge drinking. In particular, pre-existing vulnerabilities in the inhibitory control system have been associated with alcohol initiation and other risk-taking behaviors among adolescents (Norman et al. 2011; Whelan et al. 2012; Wetherill et al. 2013; Cheetham et al. 2014; Heitzeg et al. 2014). To minimize the potential effect of pre-existing vulnerabilities, subjects in the current study, binge- and non-binge drinkers were matched on various sociodemographic variables and total brain volume. Moreover, that temporal proximity of binge drinking improved model performance suggests the observed associations are not purely due to pre-existing differences.

Although the neurotoxic effects of alcohol on the developing brain have been widely reported, our study also showed a differential association of gray matter volume with respect to age cohort. Specifically, the association with binge drinking was largely attenuated for older youth. This attenuation of effects for older subjects can be clearly observed in the model fits presented in Figure 2. Our results reinforce the increase risk of negative outcomes associated with early initiation of alcohol and in particular binge drinking.

This study has several strengths, including five yearly longitudinal assessments in a cohort sequential design, enabling ascertainment of age cohort, temporal and dose associations between binge drinking and brain development. Moreover, the large sample size allowed for matching on key sociodemographic factors and baseline (pre-binge drinking) total brain volume while maintaining adequate statistical power to estimate effects and test for associations.

Despite these strengths, our findings should be interpreted in light of limitations. Our study did not examine behavioral data to capture the longitudinal association between binge drinking episodes and cognition, nor how any effects of binge drinking on the brain may mediate this relationship. Additionally, as with all observational studies, there may be confounders we did not include in the analyses that bias observed associations between binge drinking and gray matter trajectories. Finally, there was insufficient within-subject variation among binge drinkers (e.g., onset of binge drinking followed by cessation) to directly assess the impact of abstinence on the potential for recovery of gray matter volumes. However, the impact of abstinence on developmental trajectories may become estimable as the NCANDA study collects more yearly assessments.

In summary, we found that binge drinking during adolescence resulted in decreased gray matter volumes above and beyond the expected decreases from the normal maturational process in youth. This decrease was largely attenuated in later adolescence. Further, our novel findings that cortical gray matter volume decreases were greater in closer proximity to reported drinking episodes in a dose-response manner suggests a potential causal effect and tentatively raises the possibility that normal growth trajectories may be reinstated with alcohol abstinence. Given the implications of our findings, future studies with additional time points and increased temporal variation in binge-drinking patterns (including prolonged abstinence) swill be needed to examine any potential recovery effects.

## Acknowledgments

Data were collected from January 13, 2013 to January 15, 2019 and were based on a formal locked data release: NCANDA_PUBLIC_4Y_REDCAP_V01, NCANDA_PUBLIC_4Y_STRUCTURAL (https://dx.doi.org/10.7303/syn23524209; Sage Bionetworks Synapse) distributed to the public according to the NCANDA Data Distribution agreement: https://www.niaaa.nih.gov/ncanda-data-distribution-agreement.

---

**Supplementary Table:** Participant use of marijuana and tobacco products in the past year at each follow-up year, by age cohort.

| Baseline Age Cohort<br>(Years) | Mean number of marijuana use episodes in the past year (range, sd) |  |  |  |
| --- | --- | --- | --- | --- |
|  | Y1 | Y2 | Y3 | Y4 |
| Cohort 1 (<sup>3</sup> 12 & < 15) | 9.3 ([0,315], 53.19) | 16.77 ([0,361], 67.3) | 26.2 ([0,360], 81.4) | 45.7 ([0,364], 94.8) |
| Cohort 2 (<sup>3</sup> 15 & < 18) | 8.1 ([0,156], 25.2) | 28.1 ([0,361], 65.1) | 43.5 ([0,351], 89.13) | 60.1 ([0,364], 100.8) |
| Cohort 3 (<sup>3</sup> 18 & < 21.5) | 19.1 ([0,283], 64.5) | 31.9 ([0,365], 85.5) | 40.4 ([0,304], 83.9) | 52.3 ([0,365], 104.0) |
| Baseline Age Cohort<br>(Years) | Mean number of cigarette/tobacco product use episodes in the past year (range, sd) |  |  |  |
|  | Y1 | Y2 | Y3 | Y4 |
| Cohort 1 (<sup>3</sup> 12 & < 15) | 1.7 ([0,56], 9.5) | 148.6 ([0,5200], 879.0) | 210.2 ([0,7300], 1233.7) | 234.3 ([0,5475], 957.1) |
| Cohort 2 (<sup>3</sup> 15 & < 18) | 9.9 ([0,694], 74.2) | 6.1 ([0,300], 37.3) | 8.6 ([0,290], 36.3) | 20.6 ([0,1030], 113.2) |
| Cohort 3 (<sup>3</sup> 18 & < 21.5) | 16.1 ([0,480], 79.9) | 42.2 ([0,1114], 191.3) | 56.1 ([0,1200], 215.2) | 4.1 ([0,30], 8.4) |

